# Cholesterol promotes the formation of dimers and oligomers of the receptor tyrosine kinase ROR1

**DOI:** 10.1101/2025.06.19.660507

**Authors:** Alyssa Ward, Luis J. Baeza-Ballesteros, Ryan J. Schuck, Maria J. García-Murria, Rajan Lamichhane, Ismael Mingarro, Francisco N. Barrera

**Affiliations:** Department of Biochemistry & Cellular and Molecular Biology, University of Tennessee, Knoxville, USA; Departament de Bioquímica i Biologia Molecular, Institut Universitari en Biotecnologia i Biomedicina (BioTecMed), Facultat de Ciències Biològiques, Universitat de València, E-46100 Burjassot, Spain

## Abstract

ROR1 is a member of the receptor tyrosine kinase (RTK) family that plays a crucial role during organogenesis of bone and neural systems by regulating non-canonical Wnt signaling. Misregulation of ROR1 is additionally a causative factor for carcinogenesis in solid and liquid tumors. However, we have a poor understanding of how ROR1 activity is regulated. We employed a recently developed single-molecule method termed SiMPull-POP to study the oligomeric state of ROR1. RTK function is typically triggered by ligand binding, which promotes self-assembly of RTKs to form dimers and in some cases oligomers. However, our data indicate that ROR1 does not follow this paradigm. Instead, ROR1 forms dimers and oligomers in a process that is not affected by the presence of the ROR1 ligand Wnt5a. Additional experiments indicate that the transmembrane domain of ROR1 has a strong tendency to self-assemble, suggesting that this domain modulates ROR1 dimerization. Investigation into a regulatory mechanism for ROR1 self-assembly led to evaluation of the role of the lipid cholesterol, which plays pleiotropic roles in Wnt signaling. Cholesterol was found to promote the assembly of ROR1, and our results point to the transmembrane domain as the region where cholesterol exerts the regulatory effect. Taken together, our results indicate that ROR1 self-assembles in human cells; however, unlike other RTKs, this process is not stabilized by ligand binding but is instead facilitated by membrane cholesterol.

## Introduction

The receptor tyrosine kinase-like orphan receptor 1 (ROR1) plays a crucial role in the development of the nervous system, bone, and other tissues during embryogenesis (1). Activation of ROR1 in response to Wnt5a binding mediates β-catenin independent, also known as non-canonical, Wnt signaling promoting cell polarity, migration, and proliferation (2–5). Expression of ROR1 is high *in utero* but it is suppressed in adult tissues (6). An important exception are solid tumors, including breast, prostate, lung, gastric, and pancreatic cancers, and hematological malignancies such as leukemia and myeloma (7). Cancers expressing high levels of ROR1 exhibit aggressive growth and migration that correlates with poor patient prognosis (8,9). Despite its importance, we have a poor understanding of the structure and regulation of ROR1.

ROR1 has a pseudokinase domain (10) with poor catalytic efficiency, and therefore it is expected to signal through interaction with other receptors. The ROR family of receptor tyrosine kinases (RTK) has a second member, ROR2, which also participates in Wnt signaling. ROR1 binds to ROR2, forming a heteromeric complex that signals to promote cell migration and proliferation in chronic lymphocytic leukemia (11). In addition, ROR2 is able to self-assemble forming homomers, and this process promotes tumorigenesis in breast cancer(12). The assembly landscape of ROR receptors is more complex, as it entails additional hetero-interaction with the G protein-coupled receptor (GPCR) Frizzled (Fzd), and also formation of a membrane complex with the RTK ErbB3 in the case of ROR1 (13). Monoclonal antibodies that target the ROR1/2 heteromer and ROR2 homomer impair tumor growth (11,12), revealing the clinical potential of targeting protein-protein interactions in the ROR family. Here, we report single-molecule fluorescence data that indicate that ROR1 has the ability to self-assemble into dimers and oligomers, and that this process is ligand-independent.

Wnt signaling involves additional receptors, including LRP and members of the Fzd family. Lipid molecules often serve as allosteric ligands that modulate the activity, dynamics, and association of membrane proteins, including RTKs (14–19). Interestingly, the lipid cholesterol (Chol) regulates Wnt signaling in multiple ways. For example, Chol promotes the assembly of the Fzd7-LRP6-Dvl2 complex (20), and Chol levels also correlate with ROR2 expression (21). The Wnt receptors Fzd8 and LRP6 reduce membrane order by internalization of Chol-rich liquid ordered membrane domains (22). Additionally, knockdown of ROR2 in RAW264.7 macrophages negatively modulates Chol production (21). These observations indicate the involvement of Chol in Wnt signaling. Motivated by these findings, we investigated if Chol levels affected the formation of ROR1 homomers. Indeed, the results indicate that Chol enhances ROR1 self-assembly.

To investigate the molecular mechanism of ROR1 self-assembly, we performed a bimolecular fluorescent complementation assay with the isolated transmembrane domain (TMD) of ROR1. These studies revealed a strong tendency of the TMD to self-assemble, indicating that this domain is key for the formation of ROR1 homodimers. We additionally observed that assembly of ROR1’s TMD was impacted by changes in cellular levels of Chol. These results indicate that the effect of Chol on the TM domain of ROR1 can explain its effect in quaternary structure. Collectively, our findings indicate that ROR1 has a significant tendency to form dimers and oligomers. Our data indicate that ROR1 does not follow the typical ligand-dependent self-assembly, but it is instead promoted by the lipid Chol.

## Results

### ROR1 forms dimers and oligomers

To investigate if ROR1 is monomeric or instead it self-assembles, we employed SiMPull-POP, a single-molecule method that quantifies individual fluorescence photobleaching events (17). To implement this method, we first expressed in HEK293T cells full-length ROR1 labelled with a C-terminal GFP tag (**Fig. S1**). Membrane fractions were solubilized with the co-polymer DIBMA (di-isobutylene maleic acid) to form DIBMALPs, or DIBMA lipid particles, which contain native-like lipid composition (23). We imaged the obtained DIBMALPs by transmission electron microscopy (**Fig. S2**), and we observed that the nanoparticles had an average diameter of 35.1 ± 9.5 nm, within the expected range (23). After confirming the formation of DIBAMLPs, we performed a single-molecule pull down after immobilization of an anti-GFP antibody on microscope slides (**Fig. S3**). ROR1-GFP DIBMALPs were imaged via Total Internal Reflection Fluorescence (TIRF) microscopy. Single molecule videos were recorded to quantify photobleaching events. Under control conditions, ROR1-GFP exhibited 1 step and 2 step photobleaching events. We also observed ≥3 step photobleaching, which were binned into a single category (**Fig. S4**). We used published protocols to convert the photobleaching step data into fractions of monomer, dimer, and oligomers (with ≥3 ROR1 copies) (17). This step involved a statistical approach to correct for the immature GFP, which is not fluorescent. After the correction, the results showed a distribution of 49.2 ± 19.0% monomer, 30.7 ± 4.9% dimer, and 25.5 ± 11.4% oligomers (**Fig. 1**). These results therefore indicate that ROR1 self-assembles, as under these conditions roughly half of ROR1 molecules interact with other ROR1 copies.

**Figure 1:**
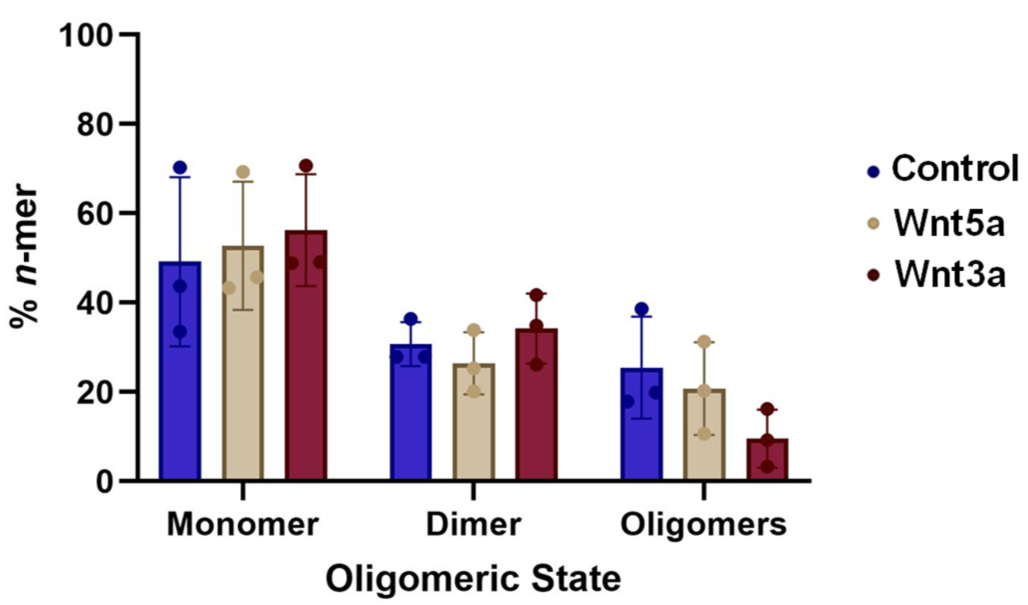
ROR1 forms dimers and oligomers. Oligomeric distribution of ROR1 isolated in DIBMALPs prepared from HEK293T cells, as determined by SiMPull-POP. N=3. Data are mean ± S.D.

Next, we studied if ligand binding affects the self-assembly of ROR1. The Wnt-family protein Wnt5a binds to both ROR1 and ROR2, promoting heterodimerization between ROR1/2 as well as homodimerization of ROR2 (11,24,25). In addition, Wnt3a binds to ROR2 (4) and promotes dimerization of other Wnt-binding receptors such as LRPs and members of the Fzd family (20). To perform this experiment, we treated the cellular membrane fractions with Wnt5a or Wnt3a, then formed DIBMALPs and finally quantified ROR1 assembly via SiMPull-POP. Strikingly, treatment with either ligand did not shift the distribution of ROR1 populations (**Fig. 1**). Overall, these results revealed that ROR1 is capable of forming membrane assemblies that are not affected by ligand addition.

### The transmembrane domain of ROR1 mediates self-assembly

We investigated next what is the region of ROR1 responsible for self-assembly. Receptor tyrosine kinases often use the transmembrane domain (TMD) to dimerize (26–28). We performed bioinformatic analysis of the dimerization of the TMD of ROR1. Both PredDimer and AlphaFold predicted a tendency for the TMD to dimerize (29,30). Indeed, two possible TMD-TMD interfaces were identified. Interface 1 is mediated by two isoleucine residues, and Interface 2 by two leucines (**Fig. S5 and Fig. S6C**). We next tested experimentally TMD’s self-assembly, and mutated the two possible interfaces.

We performed a bimolecular fluorescence complementation (BiFC) assay with the isolated TMD sequence (**Fig. 2A** and **Fig S6C**), to investigate ROR1 dimerization and the residues that mediate it (31,32). In order to identify the possible interactions, we carried out a mutational analysis aimed at blocking TMD-TMD interactions. We replaced with the small Gly residue the bulky Ile and Leu residues at Interfaces 1 and 2 (see **Fig. 2B**, with membrane insertion Δ*G*_app_ predicted as in reference (33)). We refer to these sequences as mutants 1 (Mut1) and 2 (Mut2). Additionally, we tested a scrambled sequence, and used as a second negative control the non-dimerizing H2 sequence from *E. coli* leader peptidase (Lep) (**Fig. 2B**). To perform the BiFC experiment, we transfected HEK293T cells with two plasmids bearing the two halves of a split Venus fluorescent protein (VFP), which were fused to each TMD sequence. Membrane dimerization of the TMD of ROR1 will bring together the two protein fragments, leading to assembly of fluorescent VFP (34). As a positive control and for normalization purposes, we used the TMD of glycophorin A (GpA), a hydrophobic segment that forma noncovalent homodimers within the membrane (35–39).

**Figure 2.**
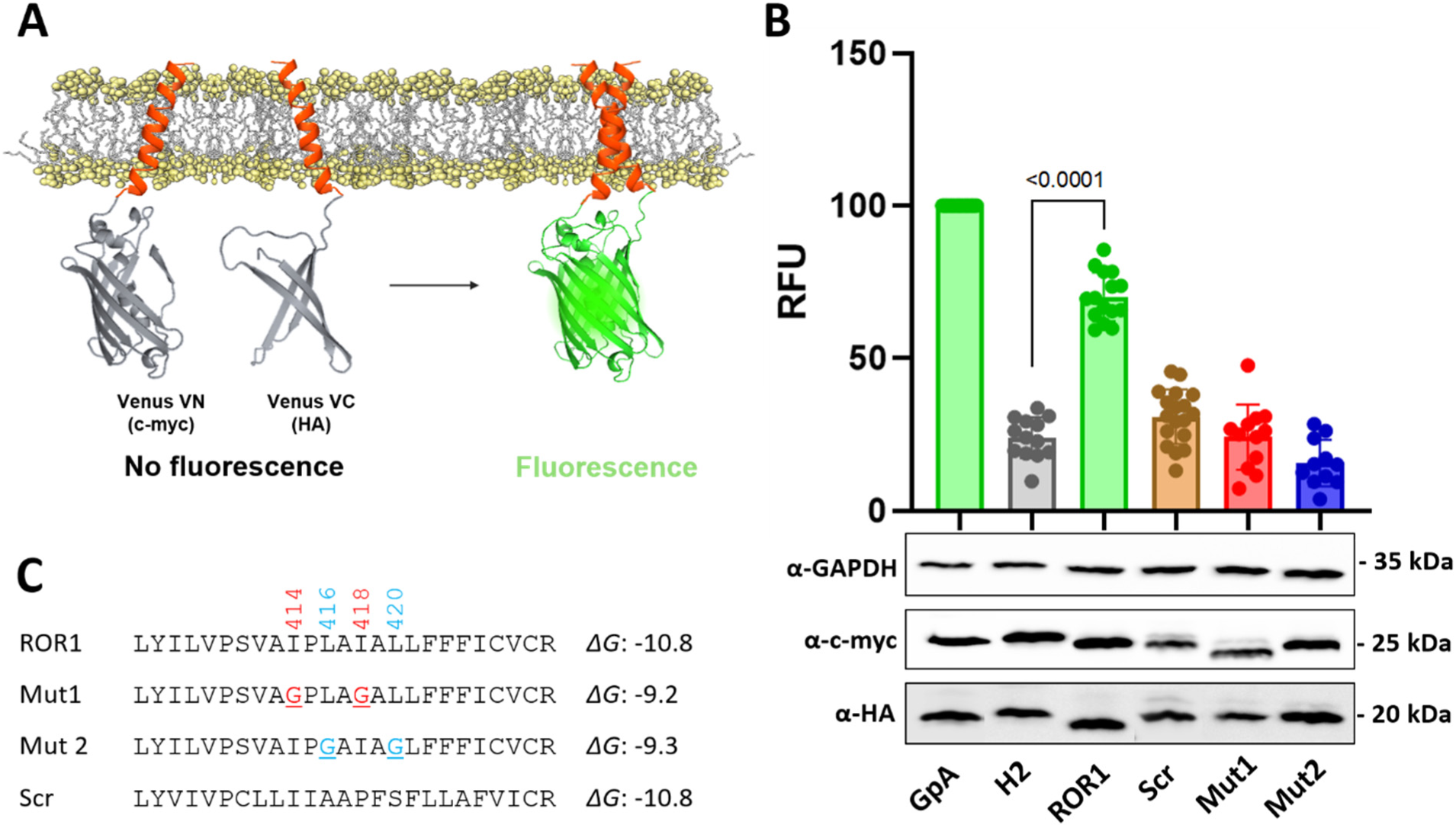
Dimerization of ROR1 TMD in HEK293T cells. **(A)** Schematic representation of the Bimolecular Fluorescent Complementation (BiFC) assay. **(B)** Amino acid sequences fused to VFP halves used in the assay and their predicted Δ*G* (Δ*G*_app_) values for membrane insertion in kcal/mol. **(C)** Relative fluorescence units (RFU) of each tested homo-oligomer in the BiFC assay. Mean and standard deviation for at least 13 independent experiments are shown. The individual value of each experiment is represented by a dot. The VN-GpA/VC-GpA homodimer is used as a positive control and to normalize values across experiments. The Lep H2 TMD was included as negative control. The homo-oligomers that produced fluorescence levels significantly higher than the H2 TMD homo-oligomer (two-tailed homoscedastic *t*-test) are highlighted in green. The lower panels display western blot data for the VN and VC constructs expression levels detected by α-c-Myc antibody and α-HA antibody respectively. An α-GPADH antibody was used as a loading control. RFU values were normalized to expression levels.

Data in **Fig. 2C** showed that the WT sequence of ROR1 TMD efficiently brought together the two VFP halves. Interestingly, mutation of both interfaces, I414G/I418G (Mut1) or L416G/L420G (Mut2), led to a severe reduction in fluorescence. In addition, the dimerization of the scrambled sequence also had low fluorescence. These results suggest specificity for dimerization of the TMD. To ensure that the results with the mutants were robust, we designed additional variants where we mutated to Ala instead of Gly. The data of the resultant Mut3 and Mut4 agreed with Gly mutation data (**Fig. S6**). We performed further controls to test the specificity of the TMD interactions, whereby we tested all pair-wise interaction possibilities between ROR1, H2, Mut1, Mut2 and the scrambled sequence. As expected, none of the combinations efficiently brought together the two VFP halves (**Fig. S7**). These data support the conclusion that the TMD of ROR1 forms a dimer or oligomer, and that this interaction occurs in a specific manner.

### Cholesterol promotes ROR1 self-assembly

Given that our data indicate that ROR1 self-assembly does not respond to ligands, we sought to investigate other cellular factors that might regulate ROR1 self-assembly. Lipid molecules can act as a ligand that modulates the activity of membrane proteins (15). Changes in the lipid composition of the plasma membrane can also alter the biophysical properties of the membrane medium, to regulate protein interactions, dynamics, and function (15). We hypothesized that Chol changes could impact the self-assembly of ROR1.

To investigate if ROR1 self-assembly in the cell is controlled by Chol, we reduced the levels of membrane Chol by acute incubation with methyl β cyclodextrin (MβCD) (40). These experiments required starvation conditions, which did not preclude self-assembly of ROR1 (compare first columns in Fig. 1 and Fig. 3B). We observed that MβCD strongly reduced Chol levels in HEK293T cells (**Fig. 3A**), as expected. Importantly, MβCD had no deleterious impact on cell viability (**Fig. S8**). We used SiMPull-POP to quantify ROR1 step populations (**Fig. S9**) and oligomerization (**Fig. 3B**) after treatment with MβCD. We observed a stark increase in the ROR1 monomer population with a subsequent decrease in the oligomer under MβCD treatment compared to control conditions. This result suggests that the high levels of Chol found in the plasma membrane promote ROR1 self-assembly.

**Figure 3:**
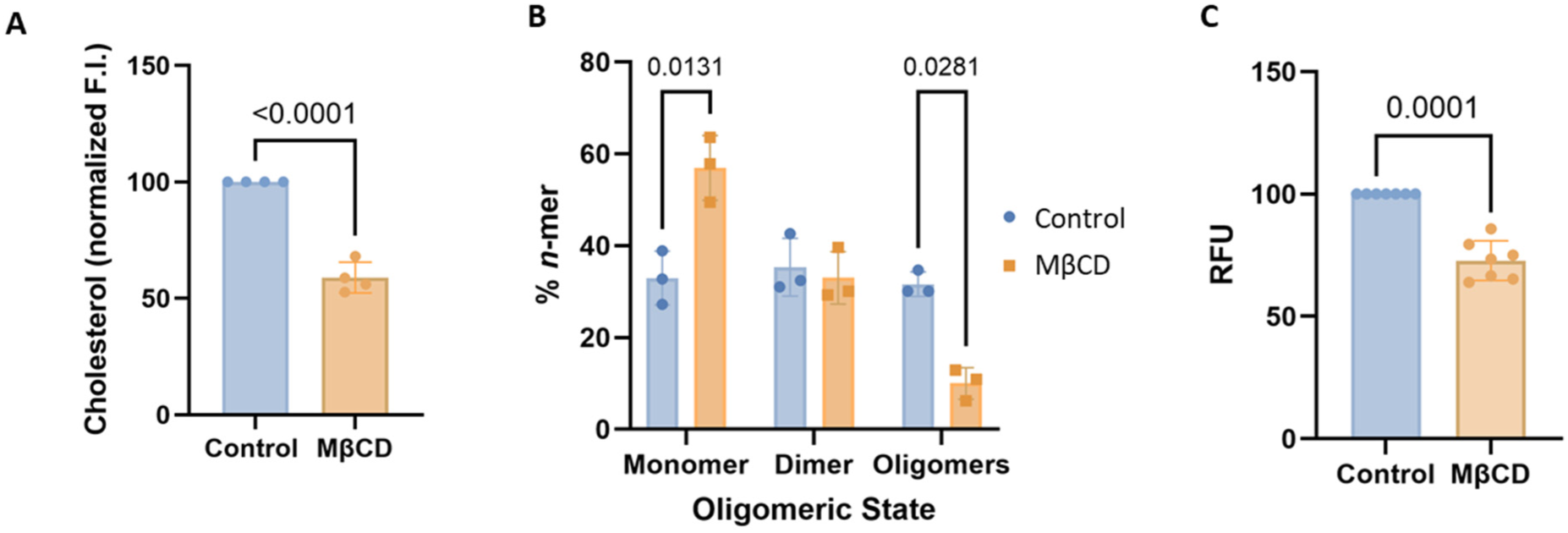
Cholesterol removal promotes monomeric ROR1. **(A)** Cholesterol levels in control conditions and after treatment with MβCD. **(B)** Oligomeric distribution of ROR1. All experiments were performed in HEK293T cells. A two-way ANOVA followed by a multiple comparison unpaired *t*-tests was run for statistical analysis. N=3. Data are mean ± S.D. **(C)** BiFC of ROR1 TMD homo-oligomers in control conditions and after MβCD treatment. Mean and standard deviation are shown. A two-tailed homoscedastic *t*-test was used for statistical analysis.

To gain mechanistic insights into the effect of Chol, we investigated whether Chol levels impact the self-assembly of the TMD. We performed the BiFC assay where the HEK293T cells were treated with MβCD to lower Chol levels. We observed that Chol removal caused a significant reduction in signal compared to ROR1 without treatment (**Fig. 3C**), indicating that a reduction of plasma membrane Chol decreases ROR1 TMD self-assembly. Taken together, the data suggest that Chol promotes ROR1 dimers and oligomers through an effect on the TMD.

To further study the possible role of Chol in ROR1 TMD interaction we examined the self-assembly of the TMD in a bacterial membrane. Bacterial membranes do not contain Chol and therefore could be used to assess ROR1 TMD dimerization in the absence of Chol (although they contain hopanoids that carry out similar functions to Chol). To investigate intramembrane contacts, we employed BLaTM, a genetic tool designed to quantitate TMD-TMD interactions in *Escherichia coli* (41,42). Briefly, the tested TMDs fused to either the N- or the C-terminal end of a split β-lactamase, βN and βC, respectively. An efficient TMD-TMD interaction facilitates the reconstitution of the complete β-lactamase and thus the growth of bacteria in selective media. The non-oligomerizing TMD of the mitochondrial protein Tomm20 (T20) was used as a negative control for membrane overcrowding and stochastic interactions. Using this approach, we tested the homo-oligomerization of ROR1 TMD. Our results indicated that ROR1 TMD has a low tendency to self-assemble (**Fig. S10**), and therefore it forms a weaker homo-oligomer in bacterial (absence of Chol) than in human cells. The bacterial results are in agreement with a previous report that applied a different dimerization assay to the isolated TMD of ROR1, which also showed a low to moderate ability to dimerize in bacterial membranes (26). Chol reduction with MβCD increases the levels of monomers in full-length ROR1 (**Fig. 3B**) as well as the isolated TMD (**Fig. 3C**). Taken together, our results support that the high levels of Chol present in the human plasma membrane promote ROR1 self-assembly.

### ROR1 Partition between L_o_ and L_d_ Domains

To investigate how Chol impacts ROR1 in the cell membrane, we studied if ROR1 accumulates in Chol-rich regions of the plasma membrane. Some Wnt receptors localize, assemble, and are internalized within liquid ordered (L_o_) regions, membrane nanodomains that are enriched in Chol (22,43,44). However, the localization of ROR1 within membrane domains is not known. We investigated ROR1 lipid domain localization using the giant plasma membrane vesicle (GPMV) assay, a well-established method to determine partitioning between L_o_ and L_d_ (liquid disordered) membrane domains (45–49). We isolated GPMVs from HeLa cells, since these GPMVs exhibit macroscopic L_o_/L_d_ phase separation at room temperature (50). Lipid phases were identified using the L_d_-specific dye Dil-C12 (51,52). Under these conditions, we could visualize the separation between the fluorescent L_d_ phase (**Fig. 4A, left**) and the non-fluorescent L_o_ phase, which correspond to the rest of the GPMV circle-like shape. We observed that ROR1-GFP was present in both phases (**Fig. 4A**), although the signal in the L_d_ phase was slightly stronger.

**Figure 4:**
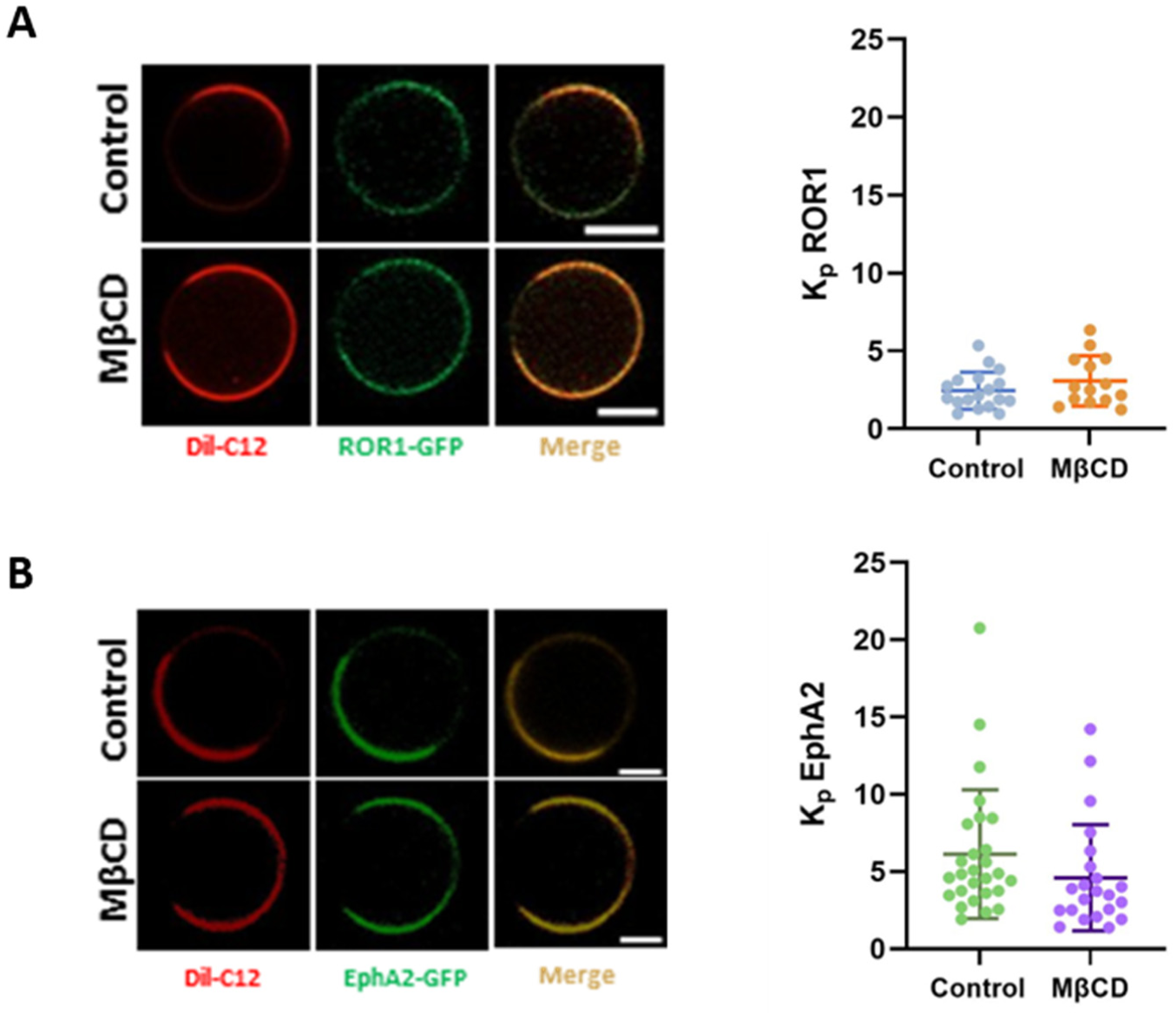
ROR1-GFP partitions into GPMV L_o_ domains more efficiently than the RTK EphA2-GFP. **(A)** Representative images of GPMVs from control and MβCD treated HeLa cells transfected with ROR1-GFP (scale bars = 5 μm) (left). L_d_ partition coefficient (*K_p_*) of ROR1-GFP in phase-separated GPMVs (right). **(B)** Representative GPMVs derived from HeLa cells transfected with EphA2-GFP in control and MβCD treated conditions (scale bars = 5 μm) (left). *K_p_* of EphA2-GFP in phase separated GPMVs (right). The *K_p_* was determined by Equation 1. Total number of GPMVs analyzed per condition were 19 (ROR1, Control), 15 (ROR1, MβCD), 28 (EphA2, Control), and 23 (EphA2, MβCD). Data are mean ± S.D., with *K_p_* values for at least 3 biological replicates.

Quantification of the average intensity in the L_d_ and L_o_ phases allows us to quantify the relative abundance of the receptor between the two phases, in the form of a partition coefficient (*K_p_*), where a *K_p_*>1 reports preference for the L_d_ phase (45). ROR1 showed a moderate preference to partition into the L_d_ phase, with *K_p_* ∼2 (**Fig. 4A, right**). As a reference, we determined the nanodomain preference of another receptor tyrosine kinase, EphA2. We observed that EphA2 has a stronger L_d_ preference (**Fig. 4B**), with a *K_p_* of ∼6 (17). These results show that ROR1 has a stronger tendency to partition into L_o_ domains that a related protein.

We investigated next if MβCD treatment affected domain partitioning. Control experiments showed that MβCD reduced similarly Chol levels in HeLa cells and that the treatment did not affect cell viability (**Fig. S11**). We incubated cells with MβCD before the treatment to induce GPMV formation. Under these conditions, however, we observed no differences in phase partitioning (**Fig. 4A**). Similar results were obtained in the case of EphA2 (**Fig. 4B**). The lack of effect of MβCD on *K_p_* might result from a balanced extraction of Chol from L_o_ and L_d_ domains. Overall, the GPMV experiments do not support preferential accumulation of ROR1 in L_o_ nanodomains (a.k.a. lipid rafts). However, comparison with EphA2 partition led us to speculate that ROR1 has a favorable interaction with Chol in GPMVs compared to similar proteins.

## Discussion

ROR1 is a prime target for therapeutic development. For example, monoclonal antibodies have recently been developed to target the ROR1/ROR2 heterodimer and ROR2 homodimer in cancer (12,53). However, ROR1 homodimerization, which is a potential alternative therapeutic target, has not been reported in the bibliography to the best of our knowledge. Here we employed the recently developed SiMPull-POP method to investigate the self-assembly state of ROR1 in the native-like environment provided by DIBMALPs. The data indicate that ROR1 populates similar levels of monomer, dimer and oligomers. ROR1/2 heterodimerization and ROR2 homodimerization are promoted by the Wnt5a ligand (11,25). We studied if ligand binding promotes ROR1 self-assembly, following the canonical activation mechanism of RTKs. Intriguingly, when we stimulated ROR1 with the ligand Wnt5a (3,4), we observed no effect, suggesting that ROR1 self-assembly is a not triggered by ligand binding. Control experiments with the ROR2 ligand Wnt3a similarly showed no change.

We investigated the molecular mechanism of ROR1 self-assembly, and our data indicate that the TMD of the receptor mediates ROR1 homodimerization. Homodimerization of other RTKs, such as the epidermal growth factor receptor (EGFR) and EphA2, is facilitated by the TMDs (26,54). Mutational experiments showed that disruption of the two predicted TMD dimerization interfaces significantly reduced self-assembly of ROR1’s TMD. Since the two interfaces are present in opposite sides of the TMD helix (**Fig. S6C**), the data suggest the existence of two types of TMD interfaces. This finding may suggest that the two predicted TMD interfaces could be engaged in alternate conformations of the ROR1 intracellular domain, as it has been observed for EGFR (55). Similarly, a recent study showcases that the TMD of ROR2 also facilitates homodimerization, and that both inactive and active dimer states were observed (11,12). Alternatively, the two TMD interfaces might be engaged at the same time in a TMD oligomer.

The lipid environment and membrane localization of RTKs can also modulate their oligomerization. For example, PI(4,5)P_2_ (phosphatidylinositol 4,5-bisphosphate) promotes the formation of homo- and hetero-dimers of the RTKs EGFR and EphA2 (14,17,56). Chol also regulates the assembly of EGFR and the insulin receptor (18,19,57–59). Chol effect on receptor interactions can be through direct or indirect mechanisms by acting as a ligand or influencing membrane localization (14,60). To assess if ROR1 self-assembly is modulated by Chol, we quantified ROR1 oligomer populations as well as the ability of the TMD to self-associate. In reduced Chol conditions we observed a significant increase in the ROR1 monomer population and a subsequent decrease in oligomers, for the full-length protein as well as for the isolated TMD. These results indicate that Cho facilitates ROR1 self-assembly, in an analogous situation where protein ligands cause formation of dimers and oligomers for other RTKs.

Most transmembrane proteins, such as HA, GT46, and FcεRI:IgE, localize to L_d_ domains (48,61,62), while a minority of transmembrane proteins like LAT and the GPCR PMP22 preferentially localize to L_o_ membrane phase (63–67). Post-translation modifications, such as palmitoylation and adaptor protein interactions have also been reported to contribute to the localization of membrane associated receptors (64,68), but are absent in ROR1. Not unexpectedly, we determined that ROR1 preferentially accumulates into L_d_ domains. However, we observed significant ROR1 levels in the L_o_ membrane phase, differently to results for another RTK, EphA2. Wnt ligands, such as those that interact with ROR1, are known to be palmitoylated and targeted to L_o_ domains presenting a potential avenue for ROR1 to localize within these domains (69,70). ROR1 localization may also depend on endocytosis events for which internalization of receptors promoting either β-catenin independent or dependent Wnt pathways occurs by different mechanisms (44,71).

In conclusion, we report here that ROR1 self-assembles in a ligand-independent manner. The formation of ROR1 homomers is instead promoted by Chol, through an effect exerted on the transmembrane domain. Our data suggest allosteric lipid-mediated self-assembly of ROR1, which is different from the canonical ligand-dependent self-assembly typical in RTKs. The identification of ROR1 dimers and oligomers represent a new potential therapeutic target for cancer.

## Experimental Procedures

### Cell Culture and Transfection

HEK293T and HeLa cells lines (ATCC) were maintained in Dulbecco’s Modified Eagle’s Medium (DMEM) supplemented with glucose, 10% fetal bovine serum (FBS), and 100 U/mL penicillin/streptomycin (1% total volume) at 37°C and 5% CO_2_. Cultures were passed or seeded once at 80% confluency and were not used after their 25^th^ passage. For expression of ROR1-GFP, cells were transiently transfected with pCAG-ROR1-GFP (Vector Builder custom plasmid) using a 3:1 polyethylenimine (PEI) (1 µg/mL) to plasmid DNA (µg). A total of 10 µg of plasmid was used for transfection of cultures seeded in a 10 cm plate and 1.5 µg for 6-well plated (Corning, CytoOne) cultures. For the transfection reaction, PEI was added to Opti-MEM media (0.5 mL for 10 cm plates and 250 µL for 6-well plates), mixed, and incubated for 5 minutes. Plasmid was mixed with an equal volume of Opti-MEM media and added to the PEI media, mixed, and incubated for 20 minutes. The transfection solution was then added to the cell culture media and incubated for 24-48 hrs.

### Cholesterol Reduction in Mammalian Cells

To reduce the level of cholesterol in HEK293T and HeLa cells, cultures were treated with methyl-β-cyclodextrin (MβCD, Acros Organics) to extract cholesterol from cellular membranes. For treatment, cultures were first rinsed with PBS (11.9 mM sodium phosphate, 137 mM NaCl, 2.7 mM KCl) then a solution of MβCD (5 mM) and HEPES (25 mM) resuspended in serum free DMEM was added to seeded cell cultures (10 mL per 10 cm plate and 2 mL per well for 6-well plates). Cultures were incubated with the solution for 1 h at 37°C and 5% CO_2_, after the solution was aspirated, and cultures were rinsed with PBS.

### Confocal Microscopy

HEK293T cells were seeded into a 24-well plate with a pre-cleaned 12 mm round #1.5 glass coverslip and grown overnight in supplemented DMEM. Cells were then transfected with the ROR1-GFP construct as above and grown overnight. To fix cells, each well was washed twice with PBS^++^ and incubated for 10 minutes with 4% paraformaldehyde at 37°C and 5% CO_2_. After fixation, each well was washed twice with PBS^++^ then stained with DAPI (1:1000) at room temperature for 5 minutes in the dark, and rinse twice with PBS++. Prepared coverslips were mounted on pre-cleaned glass microscope slides with Prolong Diamond Antifade mounting media (P36965) and allowed to cure for 24 hrs in the dark prior to imaging. Images were taken on a Leica SP8 White Light Laser Confocal System.

### MTS Cytotoxicity Assay

An MTS kit (Thermo Fisher Scientific) was used to test for cytotoxicity in HEK293T and HeLa cell lines after treatment with MβCD. Cells were seeded at 2 x 10^5^/well for each condition in a clear flat bottom 96 well plate (Corning) in supplemented DMEM (10% FBS and 1% P/S) and grown overnight at 37°C and 5% CO_2_. Cells were treated with 100 µL of 5 mM MβCD/25 mM HEPES for 1 h. Control well media was replaced with serum free DMEM during the treatment time. After treatment, wells were rinsed twice with PBS^++^ (PBS supplemented with 0.1 mM CaCl_2_ and 1 mM MgCl_2_) and grown overnight in 100 µL of phenol free supplemented DMEM. Wells were treated the following day with 10 µL MTS reagent and incubated for 1.5 hrs at 37°C and 5% CO_2_. After incubation, the absorbance of each well was measured on a Biotek Cytation V microplate reader at 490 nm. Absorbance of the wells were normalized to the control.

### Cholesterol Quantification Assay

The Amplex Red cholesterol assay (Invitrogen) was used to quantify cholesterol levels from HEK293T and HeLa cell lysates. Cells were seeded at 2 x 10^6^/well in a 6-well plate, grown for 48 hrs and harvested in a detergent-free buffer (50 mM Tris-HCl, 250 mM sucrose, 250 µM CaCl_2_, pH 7.4) and lysed by vortexing with beads (Powerbead Pro Tubes, Qiagen) for 30 minutes at 4°C. Quantification was carried out according to the manufacturers protocol and as detailed in reference (17).

### SiMPull-POP (DIBMALP preparation, smTIRF, and data analysis)

ROR1-GFP DIBMALPs were generated, imaged, and analyzed as detailed in (17). In brief, HEK293T cells were seeded into 4 x 10 cm plates (1 x 10^6^ cells) per condition, grown overnight, transfected as above, and harvested in detergent-free buffer by scraping followed by lysis via syringe passages. Membrane fractions were extracted from cell lysates by ultracentrifugation and solubilized with the co-polymer DIBMA (0.15%) overnight (shaking). Insolubilized material was separated from solubilized membrane portions (containing DIBMALPs (DIBMA lipid particles)) by ultracentrifugation. For samples treated with ligand, membrane fractions were first incubated on ice with Wnt5a (R & D Systems) (0.5 µg/mL) or Wnt3a (R & D Systems) (0.2 µg/mL) for 1 h prior to solubilization with DIBMA. A GFP calibration curve prepared with purified GFP (Thermo Fisher Scientific) in a black opaque, flat-bottom 96-well plate was used to determine the relative GFP concentration in solubilized fractions. Fluorescence was recorded on a Biotek Cytation V microplate reader. ROR1-GFP DIBMALPs were immobilized via a biotinylated GFP antibody on a PEG-biotinylated/neutravidin functionalized quartz slide with a prepared solvent filled chamber (72).For smTIRF, sample slides were imaged by a custom inverted prism-based TIRF microscope with a 465 nm cable laser. Data processing and analysis was carried out with custom IDL, Python, and MATLAB scrips as detailed in reference (17).

### Transmission Electron Microscopy

To quantify DIBMALP sizes, samples were negative stained for contrast imaging by transmission electron microscopy (TEM). A sample film was prepared on a 5-6 nm thick carbon-coated 200-square mesh copper grid (Electron Microscopy Sciences). To prepare the film, the sample was allowed to adsorb to the grid for 1 minute (grid wad placed on top singular drop of sample) followed by a 10 second rinse in ddH_2_O. The grid was then stained with 1% uranyl acetate prepared in methanol for 1 minute then allowed to air dry overnight. Filter paper was used to adsorb excess liquid from the grid between each step. Images of the prepared grids were acquired by a JEOL JEM 1400-Flash TEM (JEOL USA Inc.) with a Fischione 2400 Dual Axis Tomography Holder and Gatan OneView CMOS sensor camera. An acceleration voltage of 120 kV was used for image acquisition. DIBMALP diameters were quantified by measurements acquired in ImageJ.

### Bimolecular fluorescence complementation (BiFC)

The ROR1 TMD and the scramble sequence were cloned at the C-terminal end of the Venus VN-terminal (1-155, I152L) and VC-terminal (155-238, A206K). Mutations of BiFC TMDs were obtained using PfuPlus! DNA Polymerase mutagenesis protocol (EURx). HEK293T were grown in DMEM (Gibco) supplemented with 10% FBS in 12-well plates at 37°C, 5% CO_2_ containing 2 x 10^6^ cells/plate. After 24 h of growing, 1 µg of each plasmid encoding VN and VC were transfected using 4 µL of 1 µg/mL PEI (Merck) for each µg of DNA. Forty ng of a plasmid encoding Renilla luciferase under CMV promoter (pRL-CMV, Promega) was co-transfected for normalization purposes. Renilla were measured using the Renilla Luciferase Fash Assay Kit (Thermo), according to manufacturer’s instructions. Measurements of luminescence and fluorescence were performed 24 h post-transfection using a Multimode Plate Reader Victor X3 (Perkin Elmer). Immuno-identification of the samples was done using α-c-Myc rabbit antibody (Sigma) and followed by a secondary HRP-conjugated α-rabbit antibody (Bio-Rad) for the Venus VN quimeric protein and α-HA mouse antibody (BioLegend, Inc.) followed by a secondary HRP-conjugated α-mouse antibody (Santa Cruz Antibodies). α-GAPDH mouse antibody (Santa Cruz Antibodies) was used as housekeeping protein. Chemi-luminescence was visualized by an ImageQuant LAS 4000 (GE Healthcare).

### Cholesterol Reduction for BiFC assay

In BiFC assays, the reconstitution of the Venus protein is measured; therefore, it is essential to treat cells prior to TMD interaction, as any pre-existing interaction between the Venus protein halves would likely prevent their subsequent dissociation. To reduce cholesterol levels in HEK293T, cultures were treated with methyl-β-cyclodextrin (MβCD; Acros Organics). For this treatment, a solution of MβCD (5 mM) and HEPES (25 mM) prepared in serum-free DMEM was added to the seeded cell cultures (2 mL per well in a 6-well plate). Cultures were incubated with this solution for 1 hour at 37°C with 5% CO₂. Following incubation, the solution was aspirated, and cultures were replenished with serum-free DMEM and maintained overnight at 37°C with 5% CO₂. Luminescence and fluorescence measurements were conducted 24 hours post-transfection.

### BLaTM assay

Competent *E. coli* BL21-DE3 cells were co-transformed with N-BLa and C-BLa plasmids, version (5), containing a given TMD pair and grown overnight at 37 °C on LB-agar plates containing 34 μg/mL of chloramphenicol (Clor) and 35 μg/mL of kanamycin (Kan) for plasmid inheritance. After overnight incubation at 37 °C, overnight cultures were conducted by inoculating 5 mL of LB-medium (Cm, Kan) with 10 colonies from one agar plate, followed by overnight incubation in an orbital incubator at 37 °C, 180 rpm. An expression culture was started with a 1:10 dilution of the overnight culture in 4 mL expression medium: LB-medium (Cm, Kan) containing 1.33 mM arabinose. After 4 h at 37 °C, the expression cultures were diluted to an OD_600_ = 0.1 in expression medium. To expose the bacteria to different ampicillin concentrations, an LD_50_ culture was prepared by pipetting 100 μL of the diluted expression culture into each cavity of a 96-deep well plate (96 square well, 2 mL, VWR) containing 400 μL of expression media (final OD_600_ = 0.02) with an ampicillin gradient ranging from 0 to 300 μg/mL. The plates were incubated for 16 h at 37 °C and 250 rpm on a shaker (shaking amplitude 10 mm, KS 260 Basic, IKA) containing tips in every well to ensure a proper agitation. Cell density was measured via absorbance at 544 nm in a microplate reader (Victor X3, Perkin Elmer).

### Giant Plasma Membrane Vesicle generation and imaging

For the generation of Giant Plasma Membrane Vesicles (GPMVs), HeLa cells were seeded (2 x 10^6^/well) in a 6-well culture plate and grown overnight in supplemented DMEM at 37°C and 5% CO_2_. Cells were then transiently transfected with pCAG-ROR1-GFP plasmid (1.5 µg) as detailed above and grown overnight. Selected wells were treated with MβCD or serum free DMEM (for control condition) for 1 h at 37°C and 5% CO_2_. Wells were washed once with PBS and stored in the dark for 5-10 minutes to acclimate to room temperature followed by two rinses with buffer 1 (10 mM HEPES, 150 mM NaCl, 2 mM CaCl_2_, pH 7.4). To label L_d_ membrane phases cells were incubated with 350 pM DilC12(3) dye (Thermo Fisher Scientific) for 15 minutes in the dark at room temperature. Cells were washed twice with buffer 1 to remove excess dye then incubated with 1 mL of buffer 2 (10 mM HEPES, 150 mM NaCl, 2 mM CaCl_2_, pH 7.4, 2 mM DTT, and 24 mM (0.072%) PFA) for 1.5 hrs at 37°C and 5% CO_2_. After incubation, 100 µL of GPMV-containing media from each condition was added to a CELL view microscope slide (Greiner Bio-one) well and allowed to settle for 30 minutes. GPMVs were imaged on an inverted Zeiss LSM 580 900 Airyscan laser scanning confocal microscope (ZEISS) with a 63X oil immersion objective. Images were analyzed in ImageJ.

### Quantification of receptor membrane partitioning

For quantification of ROR1 localization, a 5-pt circle ROI outlining the Dil-C12 fluorescence/non-fluorescent circumference was first set then applied to the ROR1-GFP channel. Peak intensity graphs of the intensity vs ROI length were exported using the Multi Plot function in ImageJ for each channel. Next, the fluorescent intensities (FI) of the Dil-C12 ROI were categorized into regions of high fluorescence or low fluorescence based on a chosen FI threshold for each GPMV. Values above the threshold were categorized as L_d_ and those below were assigned as L_o_. Assignment of L_d_ and L_o_ intensities in the Dil-C12 channel were correlated with their distance coordinates/length measurement (µm) that could then be used to identify the corresponding L_d_ and L_o_ regions in the ROR1-GFP channel. The FI of ROR1-GFP within the assigned L_d_ and L_o_ regions were averaged and the partitioning coefficient (*K_p_*) of the receptor in L_d_ domains was determined as in Equation 1. For EphA2 analysis, line ROI(s) were drawn from the Dil-C12 fluorescent portion of the GPMV to the non-fluorescent side. These ROI(s) were then used to measure the peak intensities in the EphA2-GFP channel that were then exported to determine the K_p_. The peak maxima observed in the L_d_ and L_o_ regions of the EphA2-GFP channel were used to calculate the K_p_ (Equation 1) for each GPMV.

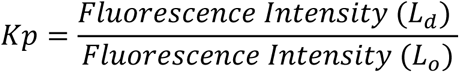

## Acknowledgements

This work was supported by NIH grants R35GM140846 (F.N.B.), R35GM142946 (R.L.), PID2023-152568NB-I00 from the Spanish Ministry of Science, Innovation and Universities (MCIN/ AEI/10.13039/501100011033) and CIPROM/2022/062 from the Generalitat Valenciana (to I.M.). We are grateful to Amit Joshi (University of Tennessee) for the use of his Zeiss confocal microscope.

