## Supplementary Figures S1-S11 for "Cholesterol promotes the formation of dimers and oligomers of the receptor tyrosine kinase ROR1"

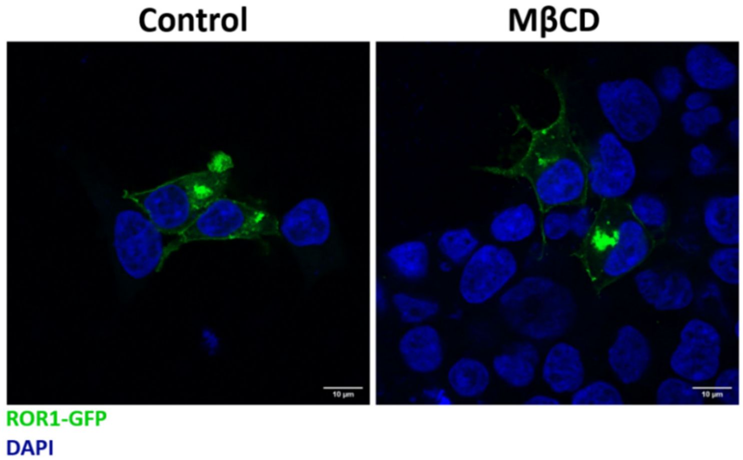


**Figure S1:** Confocal microscopy of ROR1-GFP expressed in HEK293T cells. Expression of the ROR1-GFP under control (*left*) and MβCD (*right*) conditions. Cell nuclei were stained with DAPI. Scale bars represent 10 µm.

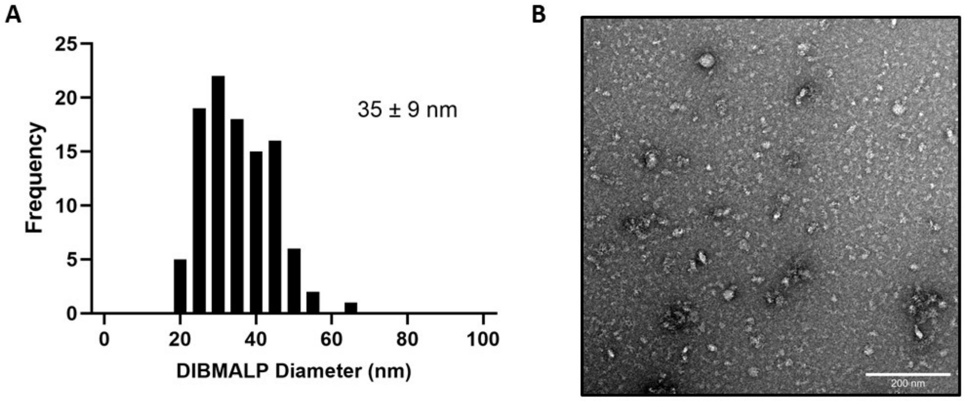


**Figure S2:** Quantification of ROR1 DIBMALP diameter by transmission electron microscopy (TEM). DIBMALPs isolated from HEK293T cells transfected with ROR1-GFP were imaged by TEM. Nanodisc diameter was quantified to be 35 ± 9 nm (mean ± S.D.). **(A)** Population distribution of DIBMALPs. **(B)** Representative TEM image of DIBMALPs. Images were collected and quantified from three biological replicates.

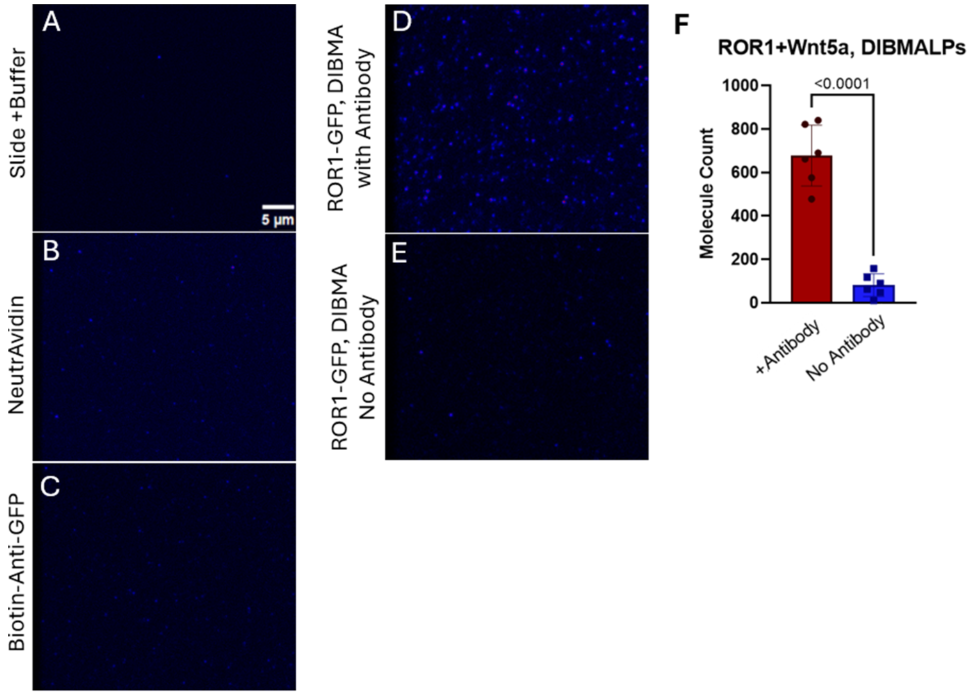


**Figure S3:** Single molecule pulldown TIRF assay of ROR1-GFP solubilized in DIBMA. **(A-C)** Representative TIRF images of a prepared slide after rinse with buffer, addition of NeutrAvidin, and incubation with the biotinylated GFP antibody. **(D-F)** Quantification of the number of molecules counted per 5 recorded TIRF movies of slides incubated with ROR1-GFP DIBMALPs, treated with Wnt5a, and after incubation with or without the biotinylated GFP antibody on the slide (representative images are shown in **(D, E)**). An unpaired *t*-tests was run for statistical analysis.

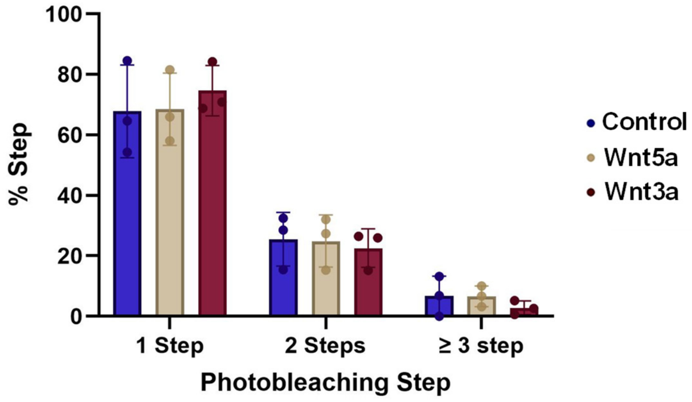


**Figure S4:** Stimulation by Wnt5a or Wnt3a does not affect ROR1 self-assembly. Percent of 1 step, 2 step, and ≥3 step populations of ROR1 isolated in DIBMALPs prepared from HEK293T cells and treated with Wnt5a (0.5 µg/mL) or Wnt3a (0.2 µg/mL). A two-way ANOVA followed by a multiple comparison unpaired *t*-tests was run for statistical analysis and showed no significance.

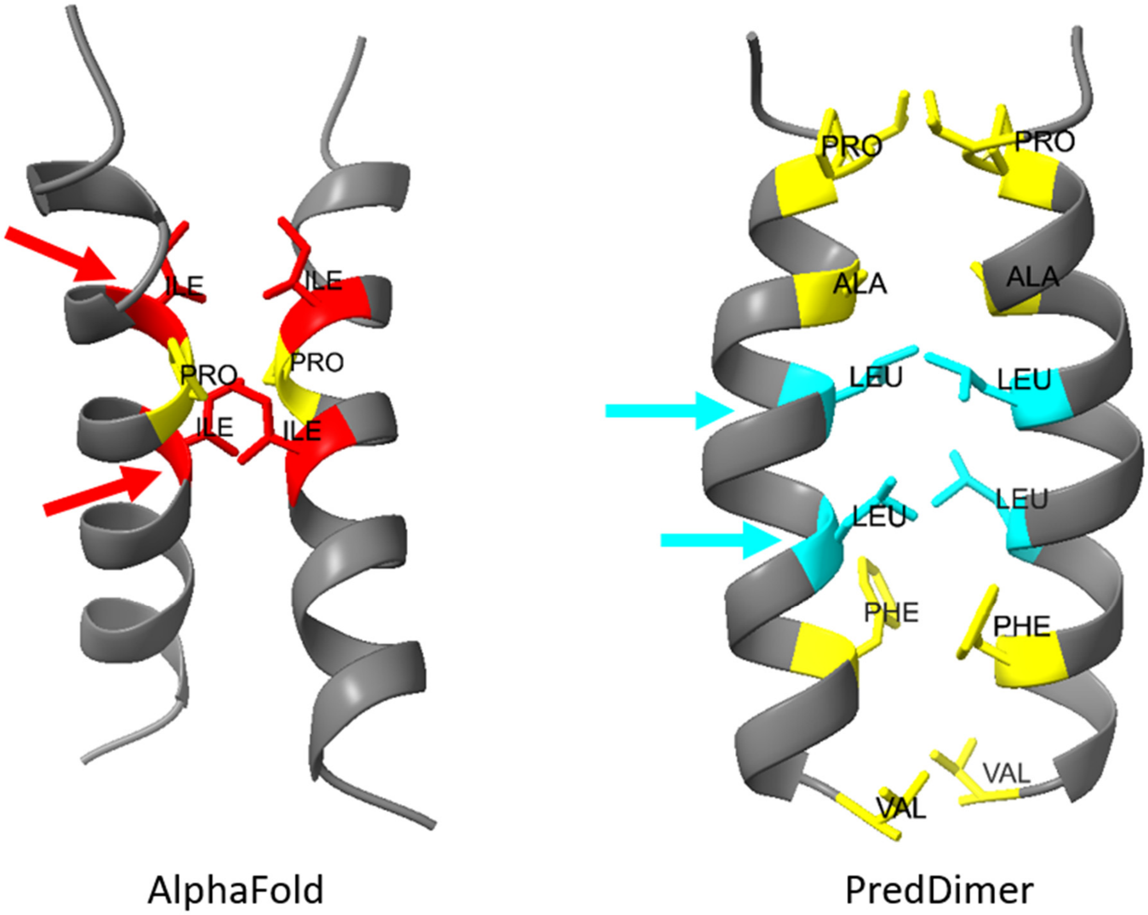


**Figure S5: ROR1 TMD Predicted Dimerization Interfaces.** (**Left**) AlphaFold predicted contact sites between ROR1 TMD helices. Highlighted residues represent predicted contact sites within 3 Å. Residues highlighted in red (Ile 414 and Ile 418) were mutated in Mut 1. (**Right**) Dimerization interface predicted by PredDimer (F-Score: 2.334). Residues represented in cyan (Leu 416 and Leu 420) were mutated in Mut 2. Residues highlighted in yellow represent additional residues predicted to form contacts or located at the TMD dimer interface. Arrows indicate location of Mut 1 and Mut 2 amino acid-pairs substitution locations. Sequence used for bioinformatic analysis: ILVPSVAIPLAIALLFFFICV.

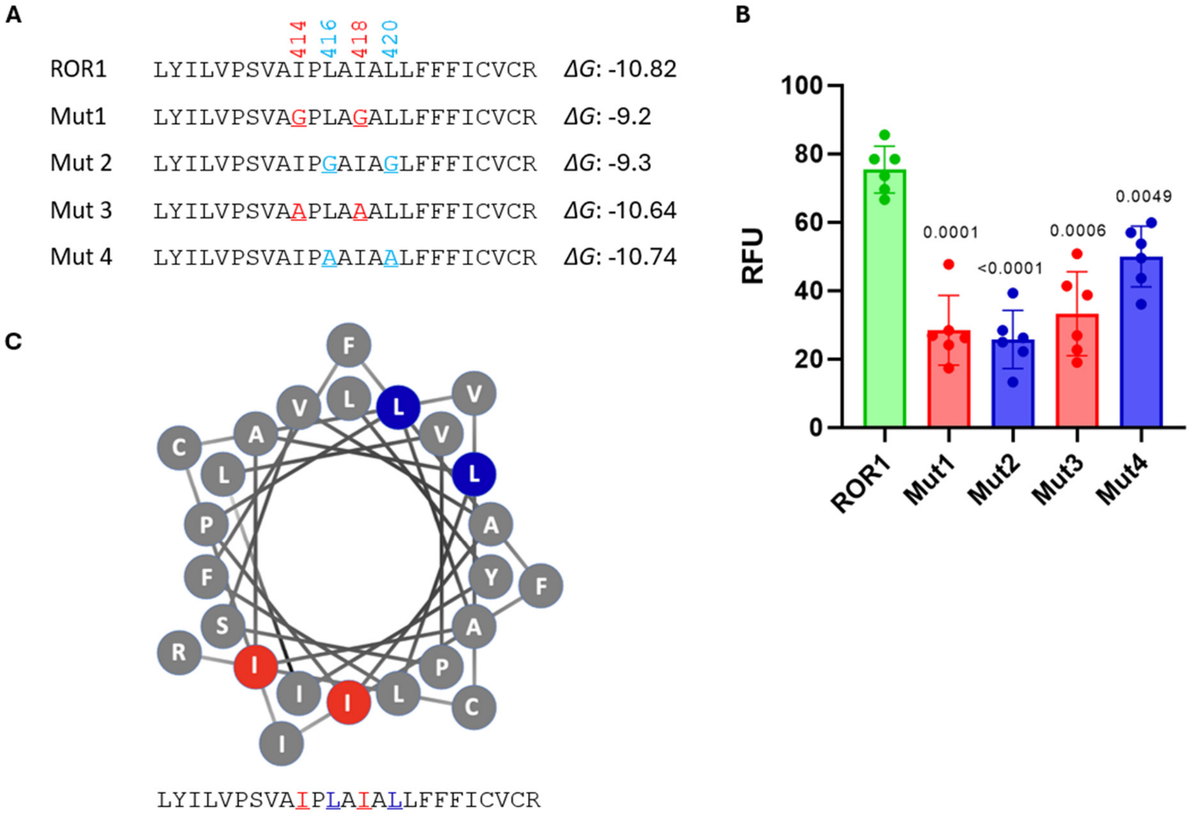


**Figure S6: Mutation of dimerization interfaces of the ROR1 TMD in HEK293T cells. (A)** Amino acid sequences of wild-type and mutant ROR1 TMDs fused to VFP halves for the BiFC assay and their predicted Δ*G* (Δ$G_{app}$) values for membrane insertion in kcal/mol. **(B)** Relative fluorescence units (RFU) of VFP fused TMD ROR1 wild-type and mutants. Mean and standard deviation are shown with each individual experiment represented by a dot. **(C)** Helical wheel projection of ROR1 TM helix highlighting in red and blue the mutated Ile- and Leu-pairs, respectively

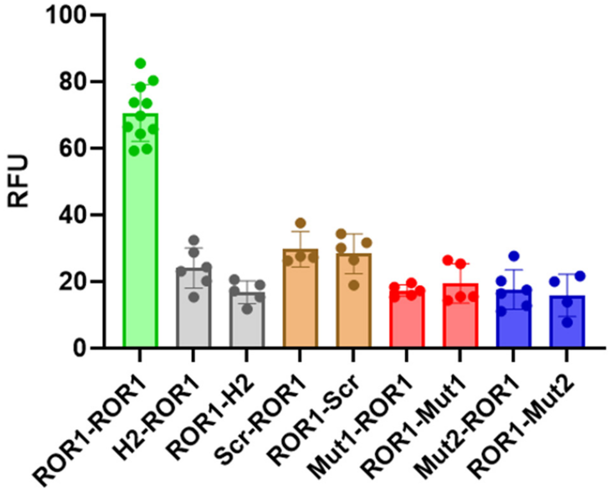


**Figure S7: Measurement of the interaction between ROR1 and the rest of the TMDs.** Relative fluorescence units (RFU) of each homo-oligomer tested in the BiFC assay. ROR1 homo-oligomerizes, as it shows levels significantly higher than the other homo-oligomers (two-tailed homoscedastic *t*-test) is highlighted in green. The rest of heterodimers are colored with different colors for each pair of TM segments in both combinations. The mean and standard deviation of at least 4 independent experiments are shown. The individual value of each experiment is represented by a solid dot.

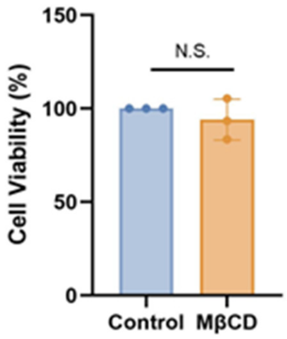


**Figure S8:** Cholesterol removal does not affect cell viability. MTS assay quantification of HEK293T cells treated with and without MβCD. An unpaired *t*-test was used for statistical analysis.

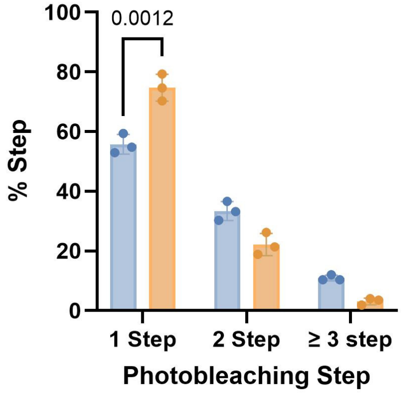


**Figure S9:** Percent step analysis of SiMPull-POP results for ROR1 isolated in DIBMALPs from HEK293T cells in control and MβCD treated conditions. A two-way ANOVA followed by a multiple comparison unpaired *t*-tests was run for statistical analysis.

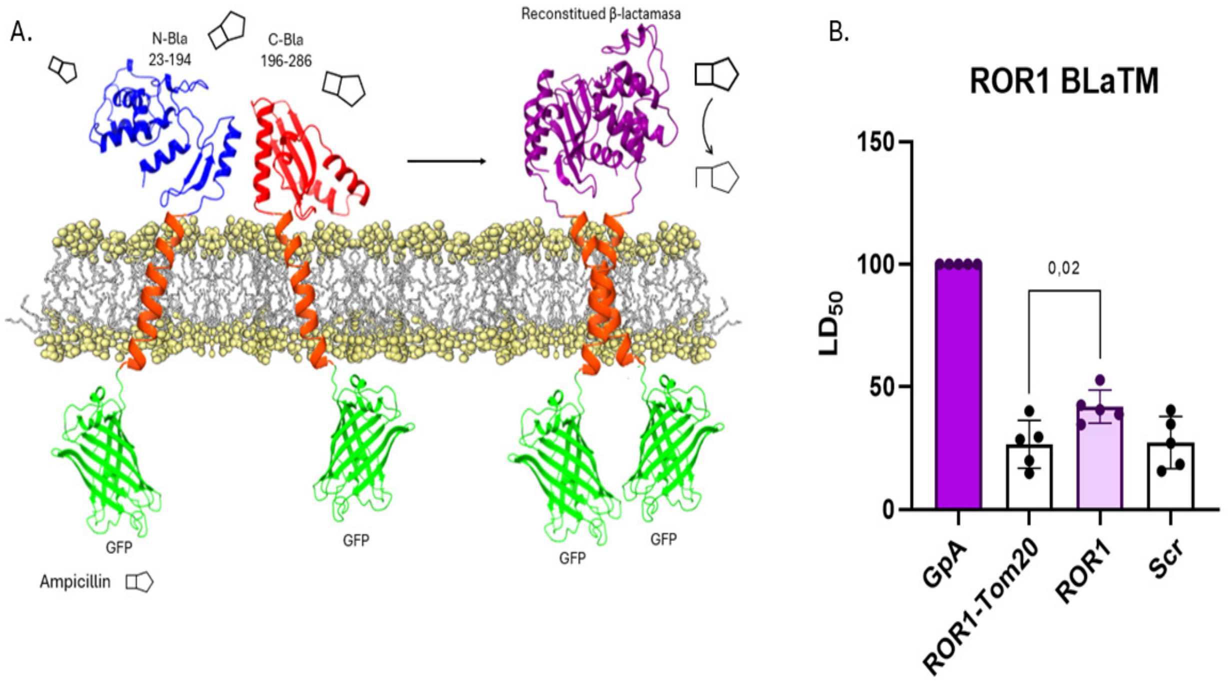


**Figure S10: ROR1 TMD interactions in bacterial membranes**. **(A)** Schematic representation of the β-Lactamase complementation (BLaTM) assay. **(B)** The βN and βC chimeras bearing the TMD of the indicated proteins were co-expressed in *E. coli*, and the resulting ampicillin $LD_{50}$ was measured. The βN ROR1–βC T20 heterodimer was used as a negative control (white), and the βN GpA–βC GpA homodimer was used as a positive control (purple) and normalization value across experimental replicates. The normalized means and SDs of at least three independent experiments ($n \geq 5$) are shown. The individual value for each experiment is represented by a solid dot. An interaction was considered positive if the observed $LD_{50}$ was significantly higher (two-tailed homoscedastic *t*-test, P value < 0.05) than the negative control.

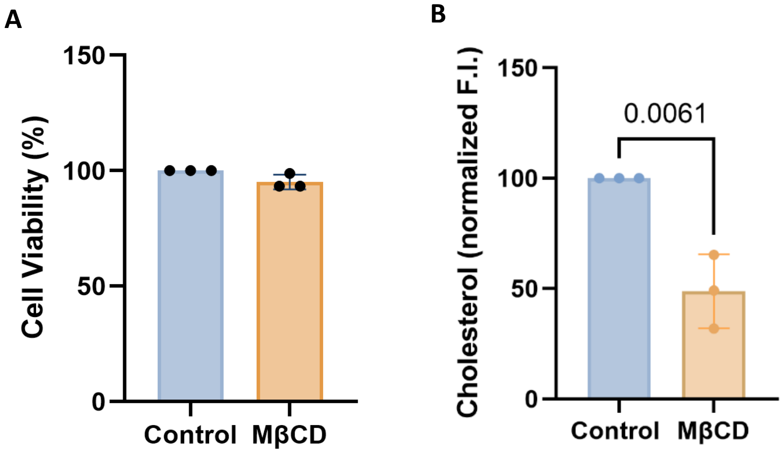


**Figure S11:** Quantification of **(A)** cell viability (MTS assay) and **(B)** cholesterol levels of HeLa cells in control and MβCD treated conditions. An unpaired *t*-tests was run for statistical analysis for both assays.
